# The role of auxin and sugar signaling in dominance inhibition of inflorescence growth by fruit load

**DOI:** 10.1101/2021.02.12.430977

**Authors:** Marc Goetz, Maia Rabinovich, Harley M. Smith

## Abstract

Dominance inhibition of shoot growth by fruit load is a major factor that regulates shoot architecture and limits yield in agriculture and horticulture crops. In annual plants, the inhibition of inflorescence growth by fruit load occurs at a late stage of inflorescence development termed the end of flowering transition. Physiological studies show that this transition is mediated by production and export of auxin from developing fruits in close proximity to the inflorescence apex. In the meristem, cessation of inflorescence growth is controlled in part by the age dependent pathway, which regulates the timing of arrest. Here, results show that the end of flowering transition is a two-step process in which the first stage is characterized by a cessation of inflorescence growth, while immature fruit continue to develop. At this stage, dominance inhibition of inflorescence growth by fruit load correlates with a selective dampening of auxin transport in the apical region of the stem. Subsequently, an increase in auxin response in the vascular tissues of the apical stem where developing fruits are attached marks the second stage for the end of flowering transition. Similar to the vegetative and floral transition, the end of flowering transition correlates with a change in sugar signaling and metabolism in the inflorescence apex. Taken together, our results suggest that during the end of flowering transition, dominance inhibition of inflorescence shoot growth by fruit load is mediated by auxin and sugar signaling.

**One-sentence summary:** Dominance inhibition of inflorescence shoot growth by fruit load is involves auxin and sugar signaling during the end of flowering transition.

## INTRODUCTION

Understanding how growing units in a shoot system are regulated, including apical and lateral buds, as well as fruits, is key to developing elite breeding lines and management tools aimed at optimizing plant architecture and increasing yield in agriculture and horticulture crops (Teichmann and Muhr, 2015; Guo et al., 2020). The activity and development of apical and lateral buds, as well as fruits, is controlled by light, temperature, hormone, sugar and nutrient signaling (Montgomery, 2008; Pfeiffer et al., 2017; Barbier et al., 2019). Moreover, these endogenous and environmental signalling pathways facilitate communication between the growing shoot apex and lateral sinks (meristems or fruits) to ensure plants adopt the appropriate architecture and reproductive capacity based on carbohydrate, nutrient and water availability (Walker and Bennett, 2018; Barbier et al., 2019).

The correlative or dominance inhibition hypothesis predicts that sinks with a high growth potential inhibit the growth of younger or subordinate organs with a lower sink activity (Bangerth, 1989; Smith and Samach, 2013; Walker and Bennett, 2018). As a result, the dominant sink is able direct water, assimilates and nutrients required for growth and development. Dominance inhibition occurs among fruits within an inflorescence or between apical and axillary buds within a shoot. Interestingly, in perennial tree crops, dominance inhibition between shoots and fruits is highly plastic, as changes in the growth potential between these competing sinks can change over the course of season. For example, a high rate of immature fruit abscission in late spring/early summer correlates with the outgrowth of preformed vegetative shoots in avocado (Salazar-García et al., 2013). Therefore, it has been hypothesized that dominance exerted by the vegetative shoot with a high growth potential results in the abscission of developing fruitlets, which have a low growth potential. However, as ‘retained’ avocado fruitlets enter a phase of rapid growth and the sink potential increases, dominance between shoots and fruits switches and shoot growth is inhibited by the developing fruit (Salazar-García et al., 1998; Ziv et al., 2014). Dominance inhibition of shoot growth by fruit load is problematic when trees maintain a high crop load, as this condition significantly reduces canopy growth resulting in a severe reduction in flowering and yield the following year (Samach and Smith, 2013; Smith and Samach, 2013). Therefore, dynamic dominance interaction between developing fruits and shoots are of significant interest, as fruit abscission and the inhibition of shoot growth by fruit load significantly reduces yield in tree crops (Samach and Smith, 2013; Smith and Samach, 2013; Sawicki et al., 2015).

Auxin is major regulator of dominance inhibition of lateral buds by the growing shoot apex, termed apical dominance (Barbier et al., 2019; Schneider et al., 2019), as well as among developing fruits within an inflorescence shoot (Bangerth et al., 2000; Smith and Samach, 2013; Walker and Bennett, 2018). In both cases, dominance inhibition is initiated by the biosynthesis and basipetal transport of auxin from the growing shoot apex or dominant fruit. For apical dominance, the polar auxin transport system (PATS) channels auxin basipetally in the stem in association with the vascular tissues (Galweiler et al., 1998). A local auxin transport system called the connective auxin transport system (CATS) also distributes this hormone in stem tissues (Bennett et al., 2016; van Rongen et al., 2019). Together, movement of auxin via the PATS and CATS indirectly inhibits bud outgrowth (Barbier et al., 2019). The canalization hypothesis predicts that a high stream of auxin channelled basipetally in the stem from the dominant shoot apex indirectly dampens auxin transport out of the lateral bud, which prevents release (Muller and Leyser, 2011). The second-messenger hypothesis reasons that high auxin concentration in the stem promotes strigolactone (SL) biosynthesis and this hormone moves into the bud to inhibit growth (Rameau et al., 2014; Barbier et al., 2019). In the bud, SL acts in part to dampen auxin transport to prevent bud outgrowth (Crawford et al., 2010; Shinohara et al., 2013). Furthermore, this mobile hormone is implicated in suppressing auxin biosynthesis and response genes in the bud (Wang et al., 2020). SL also functions to regulate key bud dormancy related transcription factors including D53/*SUPPRESSOR OF MAX2-LIKE 6*, *7* and *8* (Jiang et al., 2013; Zhou et al., 2013; Soundappan et al., 2015; Wang et al., 2015; Wang et al., 2020), as well as *BRANCHED1* (*BRC1*)/*TEOSINTE BRANCHED1* (*TB1*) (Aguilar-Martinez et al., 2007; Braun et al., 2012; Wang et al., 2020). Interestingly, studies in *Cucumis sativus* suggest that BRC1/TB1 prevents bud release in part by repressing transcription of a polar auxin transporter gene involved in branching (Shen et al., 2019). Finally, bud dormancy is maintained in part through the suppression of cell division and ribosome production (Gonzalez-Grandio et al., 2013), as well as the upregulation of abscisic acid (ABA) and jasmonic acid (JA) (Gonzalez-Grandio et al., 2017; Dong et al., 2019).

An underlying factor in dominance interaction is the ability of a developing sink to maintain a high growth potential via uptake and metabolism of sugars, including sucrose (Eveland and Jackson, 2012; Barbier et al., 2015; Pfeiffer et al., 2017). For example, the growth potential of shoot and root apices, as well as developing fruits, are dependent upon invertase activity, which functions to metabolize sucrose to glucose and fructose (Ruan et al., 2012; Bihmidine et al., 2013). In addition, sugar catabolic pathways mediated by glycolysis/the tricarboxylic acid and oxidative pentose phosphate pathway also regulate shoot growth (Wang et al., 2021). The demand of growing sinks for carbohydrates is due to the fact that sugars are key drivers of cell division and differentiation required for growth (Ruan et al., 2012; Sablowski and Carnier Dornelas, 2014). Indeed, sugar availability plays a role branching (Mason et al., 2014; Barbier et al., 2015), as wells as meristem activity (Wu et al., 2005; Pfeiffer et al., 2016). While sugars are essential for energy and cell wall biosynthesis, glucose and sucrose also function as signals that regulate plant developmental programs (Eveland and Jackson, 2012; Barbier et al., 2015), including the vegetative phase transition (Yang et al., 2013; Yu et al., 2013). In addition to sucrose and glucose, trehalose 6-phosphate (T6P) functions as a sugar signal that regulates growth in response to sucrose availability (Nunes et al., 2013; Lastdrager et al., 2014; Baena-Gonzalez and Lunn, 2020). For example, the T6P pathway regulates the vegetative and floral transition in response to the sugar availability to ensure sufficient carbohydrates are accessible to support reproductive development (Wahl et al., 2013; Ponnu et al., 2020). In addition, T6P plays a role in regulating branching and bud outgrowth in response to decapitation (Satoh-Nagasawa et al., 2006; Fichtner et al., 2017). Taken together, sugar signaling and metabolism are key drivers of plant growth and developmental processes.

In annual plants, inflorescence growth and fruit development coexist for a definite period of time before inflorescence growth ceases (Bleecker and Patterson, 1997; Nooden and Penney, 2001; Gonzalez-Suarez et al., 2020). This developmental transition is referred to as the “end of flowering” phase transition (Gonzalez-Suarez et al., 2020), which is confined to the later stage of inflorescence development (Balanza et al., 2018; Gonzalez-Suarez et al., 2020; Ware et al., 2020). Inflorescence growth cessation is mediated by fruit load, as removing these seed-bearing structures restores flower and fruit production. The inhibition of growth appears to be a separate step from senescence, which usually follows arrest (Bleecker and Patterson, 1997; Nooden and Penney, 2001; Wuest et al., 2016; Wang et al., 2020; Ware et al., 2020). A recent hypothesis predicts that inflorescence apices acquire a competency to undergo growth cessation late in inflorescence development (Ware et al., 2020). Once inflorescences acquire this competency, export of auxin from developing fruits induces growth cessation. Competency for inflorescence arrest involves the FRUITFUL (FUL)/APETALA2 (AP2) age dependent module, which indirectly regulates stem-cell homeostasis through the *WUSCHEL* (*WUS*) transcription factor (Balanza et al., 2018; Martinez-Fernandez et al., 2020). Interestingly, transcript levels for ABA signaling and response genes associated with lateral bud dormancy are higher in arrested inflorescence meristems at the end of flowering compared to active meristems during the growing phase of inflorescence development (Wuest et al., 2016). In addition, the FUL/AP1 module appears to directly regulate ABA response genes at the end of flowering transition (Martinez-Fernandez et al., 2020). Lastly, experimental studies indicate that JA may also play a role in the end of flowering phase transition (Kim et al., 2013).

Here, our results show that the end of flowering phase transition is a two-stage process that involves auxin. The first stage is marked by the selective dampening of auxin transport in the apical region of the inflorescence. Further, the transition from the first to the second stage is accompanied by an increase in auxin response in the vascular tissues where developing fruits are attached to the stem. Together, the dampening of auxin transport followed by an increase in auxin response in the apical region of the stem may function to prevent canalization required for flower production and development. Consistent with previous studies showing that sugar metabolism and signaling regulate the vegetative and flower transitions, the first stage of the end of flowering transition is associated with a significant reduction in sugar signaling and metabolism. We propose that inhibition of inflorescence growth by fruit load is regulated by auxin and sugar signaling for end of flowering transition in annual plants.

## RESULTS

### Characterization of the end of flowering phase transition

To better understand the end of flowering phase transition, the inflorescence arrest phenotype was characterized. During the growing phase of inflorescence development, the inflorescence meristem produces floral meristems, which give rise to flowers and floral organs, respectively (Fig. 1A). As flowers develop into fruits, the subtending internodes elongate, which separates the siliques. Characterization of the inflorescence arrest phenotype indicated that the end of flowering phase can be divided into two stages. During the first stage, only the apical bud, which consists of the inflorescence meristem, young unopened flower primordia and the immediate subtending internodes, transitioned to a quiescent state (Fig. 1B). In contrast, 4-6 mature flowers with developing fruits attached to elongated pedicels continued to develop (Fig. 1B). For the purposes of this study, we defined the first stage of growth cessation stage as quiescent 1 (Q1). The end of flowering phase was completed at the quiescent 2 (Q2) stage, when growth at the inflorescences apex completely ceased, including the last set of fruits to develop (Fig. 1C).

**Figure 1.**
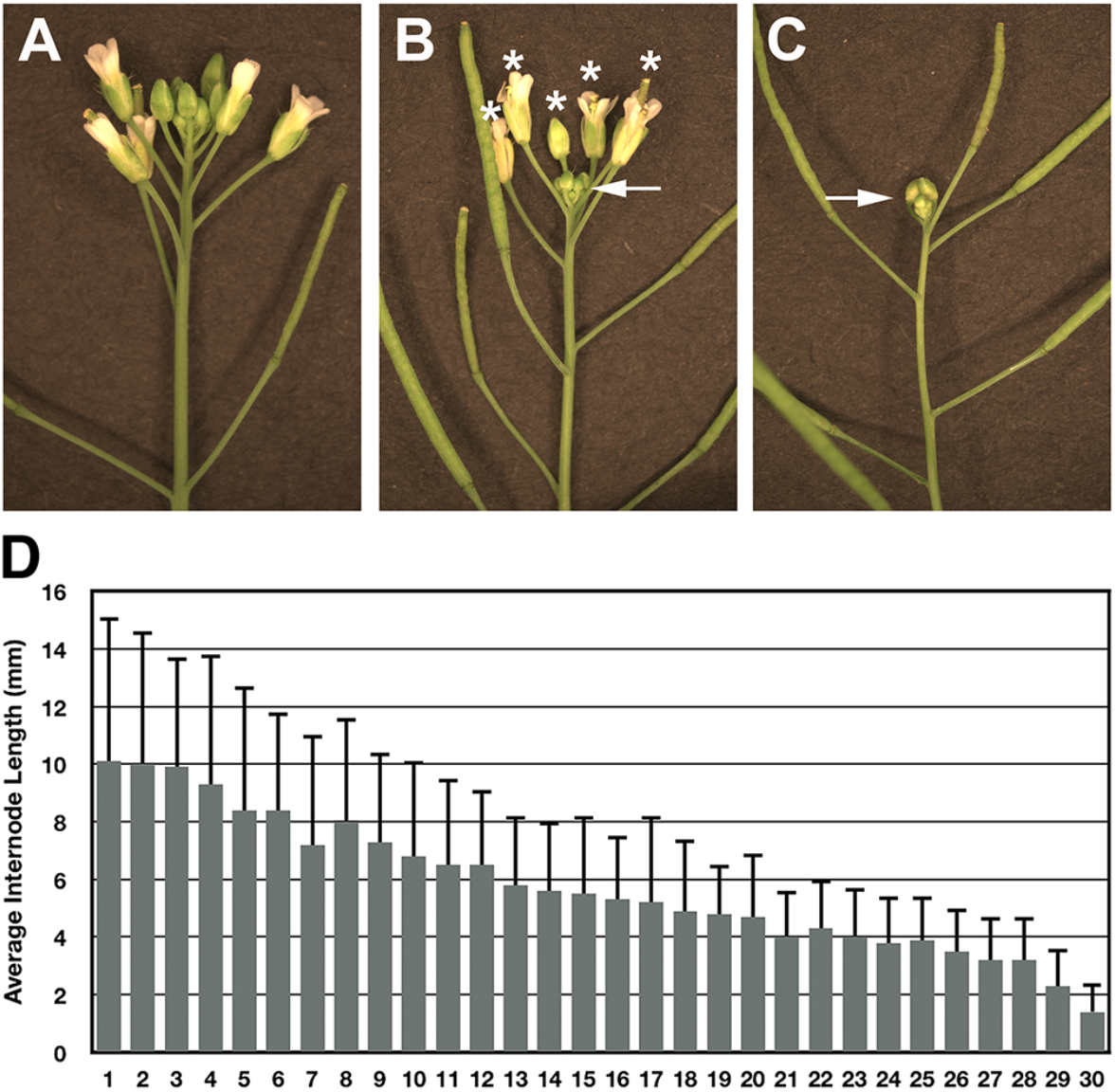
Characterization of the end of flowering transition. (A) An active inflorescence shoot apex containing numerous flowers at different stages of development. (B) Q1 inflorescence shoot apex with a white arrow pointing at the compact quiescent apex, which consists of young unopened flower buds. The asterisks mark mature flowers with developing fruit attached to elongated pedicels. (C) Image inflorescence shoot apex at the Q2 stage of development, in which growth at the apex, including the fruits, ceased. (D) Average internode length was determined for the last 30 internodes produced after the end of flowering transition. Note: internode 30 is the last internode to elongate before Q1 stage arrest.

In actively growing Arabidopsis inflorescences, the shoot meristem allocates cells that give rise to flowers and internodes (Serrano-Mislata and Sablowski, 2018). The gradual decline in meristem size indicates that meristem activity decreases during inflorescence development (Balanza et al., 2018; Wang et al., 2020). To further support this hypothesis, the average length for the last 30 internodes produced on the primary stem was determined. Results showed that over the course of inflorescence development, internode length gradually declined (Fig. 1D). The steady decline in internode development indicates that the meristem allocates fewer and fewer cells to support stem growth due to a gradual decrease in meristem activity.

### Transition to the Q1 stage is associated with a cessation of meristem activity

To evaluate the effect of fruit load on growth processes in the inflorescence apex, a series of mRNA *in situ* hybridizations were performed with genes that control meristem activity. To demonstrate that cessation of inflorescence growth occurred at the Q1 stage, the cell division marker, *CYCLIN DEPENDENT KINASE B1;1* (*CDKB1;1*) was used as a marker to assess whether the shoot apex was active (Segers et al., 1996). Results showed that *CDKB1;1* was expressed in inflorescence and flower meristems, as well as the vasculature of actively growing inflorescence apices (Fig. 2A). In contrast, transcripts for *CDKB1;1* were not readily detected in Q1 shoot apices (Fig. 2B). The *HISTONE H4* gene, which also serves as a cell division marker (Krizek, 1999; Gaudin et al., 2000), was not expressed in Q1 apices compared to active inflorescence apices (data not shown). *SHOOTMERISTEMLESS* (*STM*) is a regulator of shoot meristem identity (Long et al., 1996). Therefore, to determine if the fate of the inflorescence meristem cells had changed during Q1, the expression pattern of *STM* was examined. Results showed that *STM* was expressed in both active and the Q1 inflorescence and floral meristems (Fig. 2C and D).

**Figure 2.**
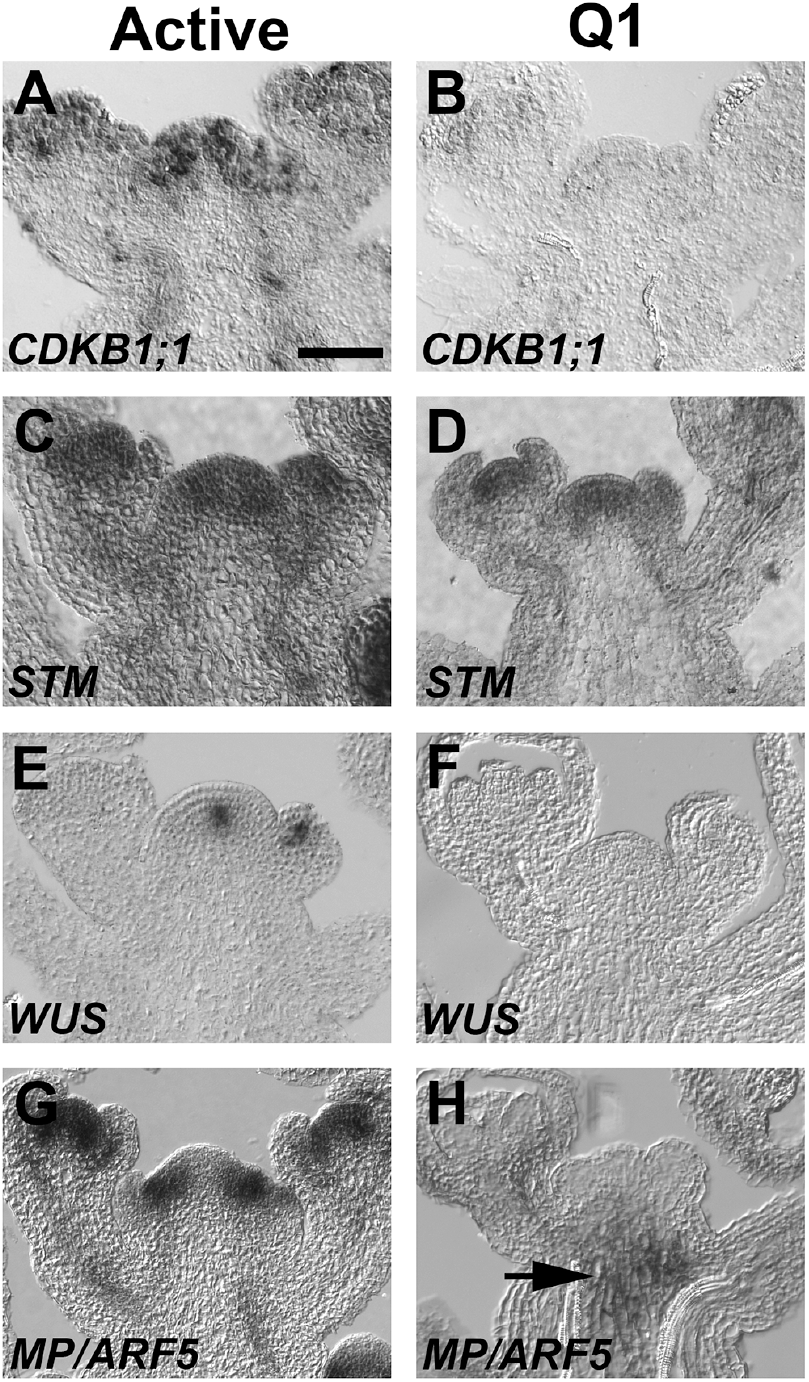
Expression patterns for key genes that control shoot growth. Gene expression patterns were shown for (A, C, E and G) active and (B, D, F and H) Q1 apices. (A and B) *CDKB1;1* is a mitotic regulator expressed in shoot meristem (Segers et al., 1996). (C and D) *STM* is a shoot meristem identity gene (Long et al., 1996). (E and F) *WUS* is key regulator of stem cell homeostasis (Laux et al., 1996). (G and H) *MP* regulates flower formation and vascular development (Hardtke and Berleth, 1998). (A) The length of the bar is 50 μm. (H) Arrow points at the *MP* expression in the sub-apical region of the meristem and pith cells in the Q1 apex.

To investigate the impact of fruit load on stem cell homeostasis, the expression pattern for *WUSCHEL* (*WUS*) was evaluated in active and Q1 inflorescence apices. In active inflorescence apices, *WUS* was expressed in the central domain of the inflorescence meristem (Fig. 2E) (Laux et al., 1996; Clark et al., 1997). In contrast to active inflorescence apices, *WUS* expression was not detected in Q1 meristems (Fig. 2F). *MONOPTEROS* (*MP*)/*AUXIN RESPONSE FACTOR* (*ARF5*) encodes an auxin response factor that is expressed in the periphery of the shoot meristem where it controls auxin mediated leaf and flower formation, as well as vascular development (Przemeck et al., 1996; Hardtke and Berleth, 1998; Schuetz et al., 2008). In active inflorescence apices *MP/ARF5* expression was detected in peripheral region of the shoot meristem and vascular tissues of the inflorescence apex (Fig. 2G). Interestingly, the expression pattern of *MP/ARF5* was altered in Q1 inflorescence apices, as the mRNA localized to the subapical region of the inflorescence meristem (Fig. 2H). Further, *MP/ARF5* expression was no longer detected in the periphery of the inflorescence meristem, as well as the quiescent floral meristems and vasculature tissues of the stem and pedicels (Fig. 2H). Taken together, the expression studies show that key determinants of cell division, stem cell homeostasis and auxin-mediated organogenesis are suppressed at the Q1 stage. However, meristem identity is maintained in Q1 meristems, as indicated by the expression of *STM*.

### Selective inhibition of auxin transport in the apical inflorescence stem correlates with arrest

We speculated that the end of flowering transition involved dominance inhibition of inflorescence growth by fruit load. Moreover, dominance inhibition was predicted to correlate with the selective inhibition of auxin transport in the apical region of the stem below the inflorescence apex. To test this hypothesis, basipetal auxin transport was measured in two sets of stem segments during inflorescence development using ^14^C-indole-3-acetic acid (^14^C-IAA). First, auxin transport was determined in apical stem (AS) segments (Fig. 3A and B), from the stem region just below the inflorescence apex to the site of stem where developing fruits were attached. The region of the stem were developing fruits are attached was referred as the zone of fruit development (ZFD; Fig. 3A and B, white box). In basal stem (BS) segments, auxin transport was also measured below the ZFD (Fig. 3A and B).

**Figure 3.**
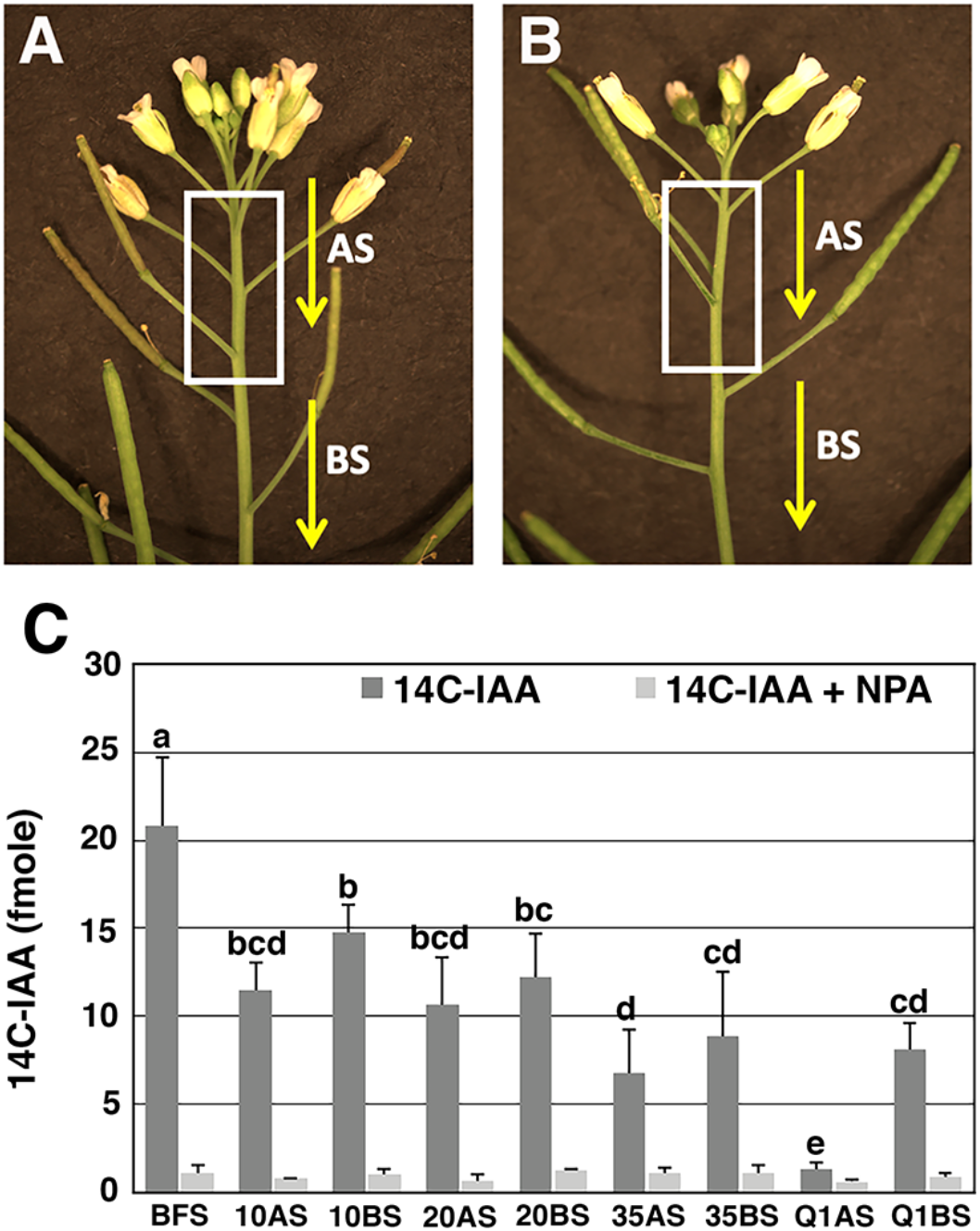
^14^C-IAA transport in stem segments during inflorescence development. Images of (A) active and (B) Q1 inflorescence shoots. The white box marks the region of the stem where developing fruits are attached. This region is referred as the zone of fruit development (ZFD). IAA transport was determined in apical stem (AS) segments and basal stem (BS) segments relative to the ZFD. (C) Radiolabeled IAA transport was measured during inflorescence development starting at a time just before the first fruit set (BFS). After fruit set, ^14^C-IAA was determined in apical stem (AS) and basal stem (BS) segments when 10, 20 and 35 fruits were produced, as well as the Q1 stage. The light colour boxes represent control stem segments in which ^14^C-IAA transport was measured in the presence of naphthyphthalamic acid (NPA), an inhibitor of polar auxin transport. The letters above the bars determine whether differences in ^14^C-IAA transport were statistically significant using analysis of variance, Tukey’s honest significant difference.

In this analysis, auxin transport was measured early in inflorescence development before fruit set (BFS) and results showed that these stem segments transported an average 20.8 fmoles ^14^C-IAA (Fig. 3C). After 10 fruit set, a significant decline in radiolabeled IAA transport occurred in AS and BS segments (Fig. 3C). The level of ^14^C-IAA transport was maintained in AS and BS segments up to the time in which 20 fruit set. At the 35-fruit set time point, an apparent further decline in radiolabeled IAA transport occurred (Fig. 3C). At the Q1 stage, auxin transport was severely reduced in AS segments, as these segments transported an average 1.3 fmoles of ^14^C-IAA (Fig. 3C). Interestingly, the average amount of ^14^C-IAA transported in Q1 BS segments was 8.1 fmoles, which was similar to the transport capacity of 35 BS segments. In addition, there was an apparent trend in which BS segments displayed higher ^14^C-IAA transport capacity than AS segments when shoots produced 10, 20 and 35 fruits (Fig. 3C). Lastly, removing fruit at the Q2 stage, restored inflorescence growth and IAA transport in AS segments (Supplemental Fig. S1). In summary, these results indicate that fruit load modulates auxin transport and supports the hypothesis that inflorescence growth arrest correlates with a selective dampening of auxin transport in the AS segments at the Q1 stage.

### Auxin response increase is primarily associated with Q2 stage of growth cessation

To further characterize the role of auxin in meristem arrest, auxin response was examined in active and arrested inflorescences at the Q1 and Q2 stages using the synthetic *DR5* auxin responsive promoter fused to the reporter gene beta-glucuronidase (GUS) (Ulmasov et al., 1997). In active inflorescence apices, *DR5:GUS* activity was detected primarily in young floral buds (Fig. 4A). In addition, *DR5:GUS* expression was also detected in fruits (Fig. 4A) and pedicels, particularly at the base of the developing fruits (Fig. 4B). At the Q1 stage, *DR5:GUS* was detected in the dormant floral buds but to a lesser extent compared to active inflorescence apices (Fig. 4C). In addition, *DR5:GUS* was detected in the last set of developing fruits (Fig. 4C). Interestingly, in approximately 37% of Q1 *DR5:GUS* inflorescences, GUS activity was also apparent in the ZFD and throughout most the pedicels of developing fruits (Fig. 4C and D). GUS activity in the remaining 63% of Q1 *DR5:GUS* inflorescences was below the level of detection similar to actively growing inflorescences (data not shown). At the Q2 stage, auxin response was detected primarily in the inflorescence stem and pedicels just below the inflorescence apex where the last set of fruit developed (Fig. 4E and F). The “stripe-like” pattern of DR5:GUS activity in the stem suggests that auxin response was induced in the vascular tissue of the main stem and pedicels (Fig. 4F). To test this hypothesis, histological experiments were performed in active and Q2 inflorescence stems below the inflorescence apex where the last 2-3 fruits had set. Results showed that *DR5:GUS* was not detected in cross sections through the ZFD during active inflorescence growth (Fig. 4G). In contrast, *DR5:GUS* activity was readily observed in vascular cells of the Q2 stems where the last set of fruit developed (Fig. 4H). Taken together, results show that auxin response increases in the vascular tissues of the apical stem, which corresponds to the site where the last set of fruit completed their developmental program during the transition from Q1 to Q2.

**Figure 4.**
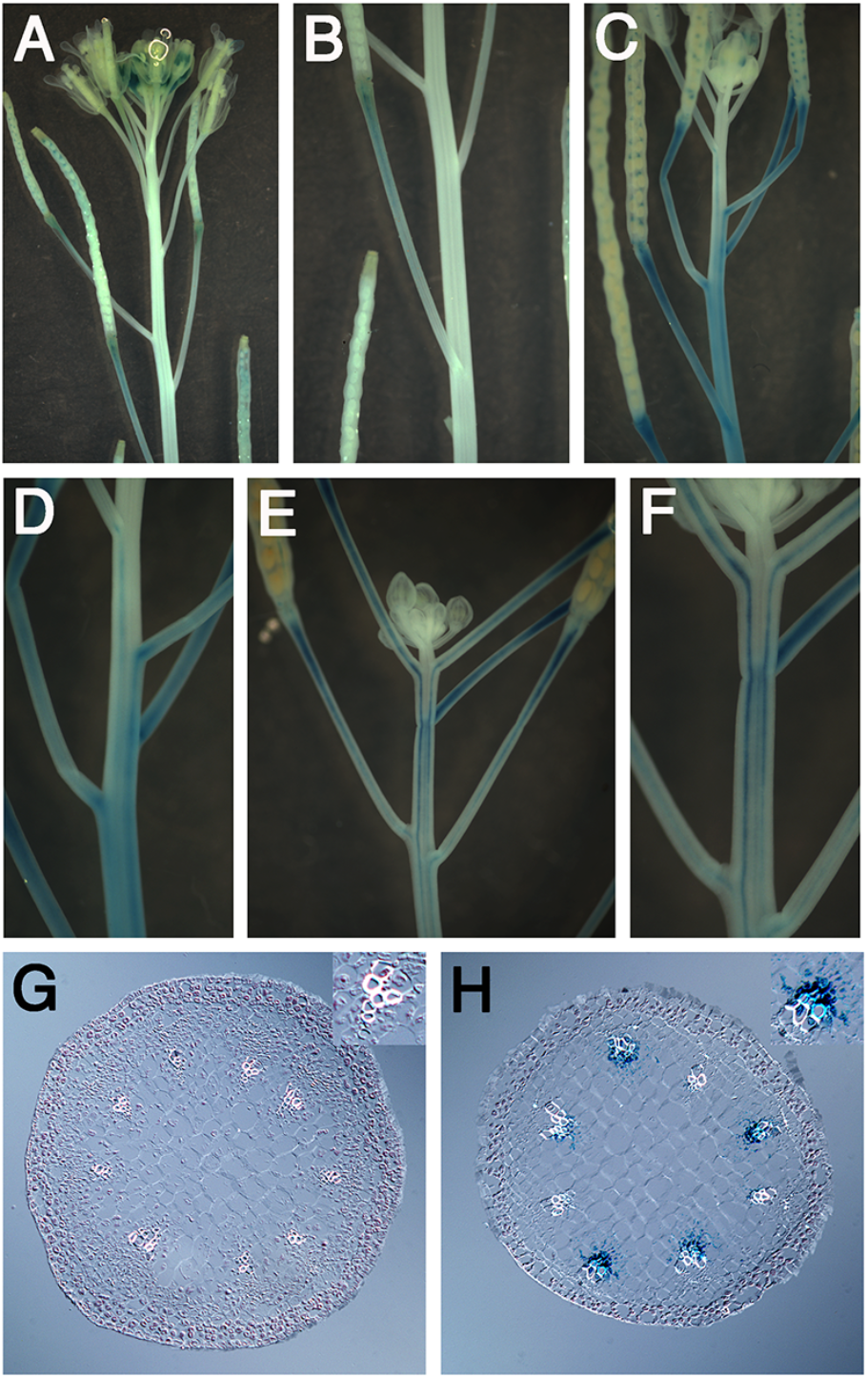
A change in auxin response in apical inflorescence stems at end of flowering transition. *DR5:GUS* staining patterns in (A) active, (C) Q1 and (E) Q2 shoot apices. Close up of stems where developing fruits are attached in (B) active, (D) Q1 and (F) Q2 shoot apices. Histological cross sections in the apical stem for an (G) active and (H) Q2 stage inflorescence. Note: inset of vascular bundle displayed in upper right corner of each image. In actively growing shoots, the section shown in (G) was through the region of the stem where the developing fruits were attached. The section displayed in (H) was in a region of the stem where the last fruits set.

### Carbohydrate status is reduced in Q1 inflorescence apices

As sugar signaling and metabolism plays a critical role in developmental phase transitions (Poethig, 2013; Wang, 2014), and influences auxin signaling and transport (Le et al., 2010; Lilley et al., 2012; Sairanen et al., 2012; Barbier et al., 2015; Lauxmann et al., 2016), we examined the carbohydrate status of inflorescence apices after the transition to the Q1 stage. The expression patterns of key genes involved in sugar signaling, metabolism and transport were investigated. *TREHALOSE 6-PHOSPHATE SYNTHASE 1* (*TPS1*) is an essential enzyme that catalyzes T6P from glucose-6-phophate and UDP-glucose (Lastdrager et al., 2014; Baena-Gonzalez and Lunn, 2020). As expression of *TPS1* correlates with T6P levels during inflorescence development (Wahl et al., 2013), this biosynthetic gene was used as a marker to assess the T6P pathway in Q1 shoot apices. Results showed that *TPS1* was primarily expressed in the vascular system of active inflorescence apices, as well as the flanks of the inflorescence meristem (Fig. 5A). In contrast to actively growing inflorescence apices, *TPS1* was not detected in the vascular system or inflorescence meristems in Q1 apices (Fig. 5B). Invertases are key enzymes involved in regulating sink activity in meristems and fruits (Ruan et al., 2012; Bihmidine et al., 2013). The *CYTOSOLIC INVERTASE 1* (*CINV1*) and related genes in *Oryza sativa* (*OsCYT-INV1*), *Lotus japonicas* (*LjINV1*) and *Solanum lycopersicum* (N16) are required for growth and carbon partitioning (Lou et al., 2007; Qi et al., 2007; Jia et al., 2008; Barratt et al., 2009; Welham et al., 2009; Barnes and Anderson, 2018; Leskow et al., 2020). In active inflorescence apices, *CINV1* was expressed in inflorescence and flower meristems, vascular cells and young floral organ primordia (Fig. 5C). Similar to the results obtained with *TPS1*, *CINV1* expression was not detected in the Q1 apices, (Fig. 5D) indicating that sucrose metabolism and sink activity is highly reduced in arrested meristems. To investigate a possible effect of inflorescence growth arrest by fruit load on carbohydrate partitioning, the *SUCROSE TRANSPORTER 2* (*SUC2*) was examined, as this transporter is expressed in the vasculature of inflorescence stems (Truernit and Sauer, 1995; Gottwald et al., 2000). Results showed that *SUC2* was expressed in the vasculature tissues of active and Q1 apices (Fig. 5E and F). While *SUC2* is expressed in Q1 apices, the decrease in *CINV1* and *TPS1* expression indicates that carbohydrate status was reduced when shoot apices transition to the Q1 stage.

**Figure 5.**
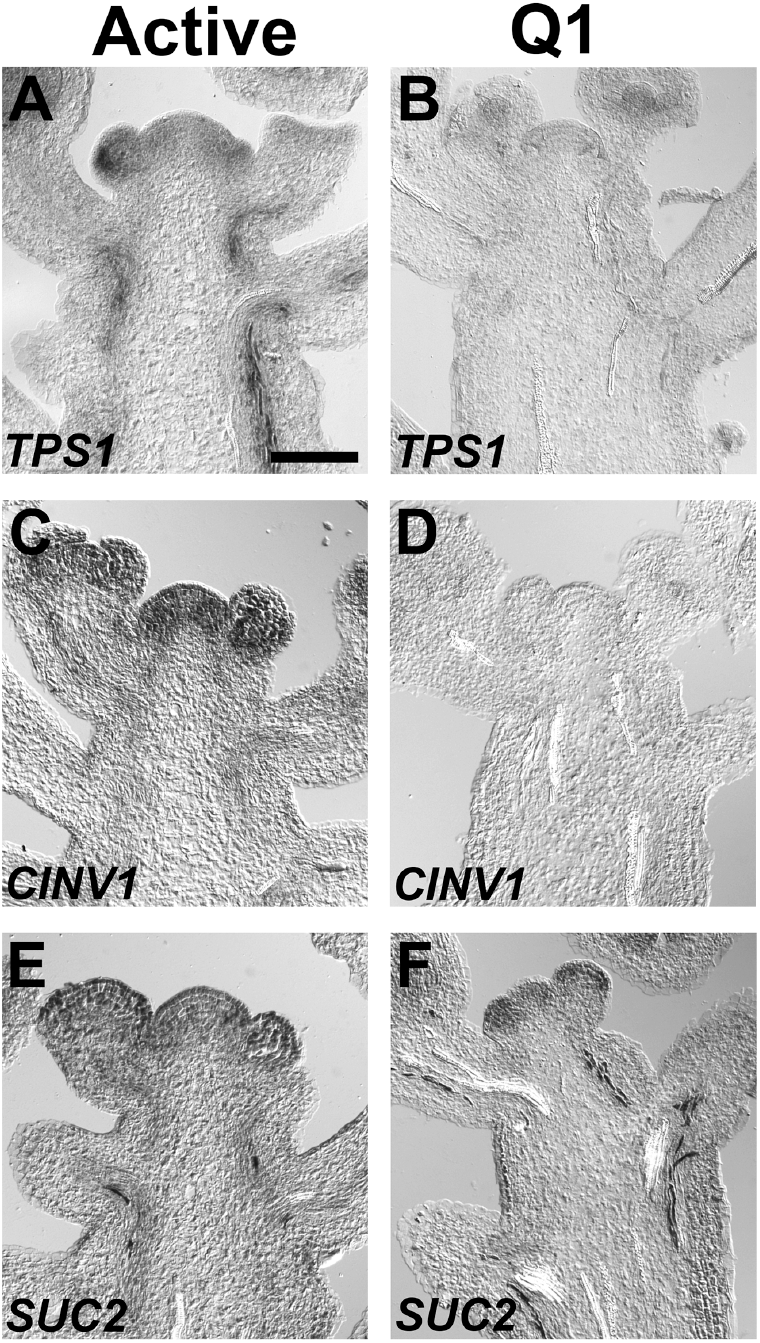
Expression patterns for sugar signaling, metabolism and transport genes. Gene expression patterns were determined in (A, C and E) active and (B, D and F) Q1 apices. (A and B) *TPS1*, which is expressed in the vasculature of the inflorescence stem, was used as a marker to assess the T6P pathway (Wahl et al., 2013; Ponnu et al., 2020). (C and D) *CINV1* is an invertase, which is required for growth (Barratt et al., 2009). (E and F) *SUC2* is a sucrose transporter expressed in the vascular tissues of inflorescences (Truernit and Sauer, 1995; Gottwald et al., 2000). The length of the bar is 50 μm.

To further investigate a role for sugar metabolism and signaling in the end of flowering phase transition, glucose, fructose and sucrose were measured in inflorescence apices before and after 15 fruit set, as well as the Q1 stage. In addition, these sugars were measured in developing fruits when 15 fruits set and at the Q1 stage. Results showed that the levels of glucose and fructose were similar in active inflorescence apices before and after 15 fruit set (Fig. 6A and B). In contrast, the levels of these monosaccharides were significantly reduced in Q1 inflorescence apices. (Fig. 6A and B). In active and Q1 inflorescence apices, the level of sucrose was similar, indicating that sucrose transport was not affected at the Q1 stage (Fig. 6C), which is consistent with the expression of *SUC2* in arrested inflorescences. Together, these results indicate that sucrose metabolism but not transport is significantly reduced when inflorescence apices transition from an active to a quiescent state. In developing fruits, the levels of glucose, fructose and sucrose were significantly higher compared to active inflorescence apices (Fig. 6A and B). These results indicate that developing fruits have a higher sink potential than active inflorescence apices during inflorescence development. Interestingly, developing fruits at Q1 had the highest levels of glucose and fructose indicating that sucrose metabolism is increased in fruits at the end of flowering phase (Fig. 6A and B). Taken together, results from above indicate that a change in sugar signaling and metabolism in arrested inflorescence apices and developing fruits is associated when active inflorescence apices transition to the Q1 stage during the end of flowering transition.

**Figure 6.**
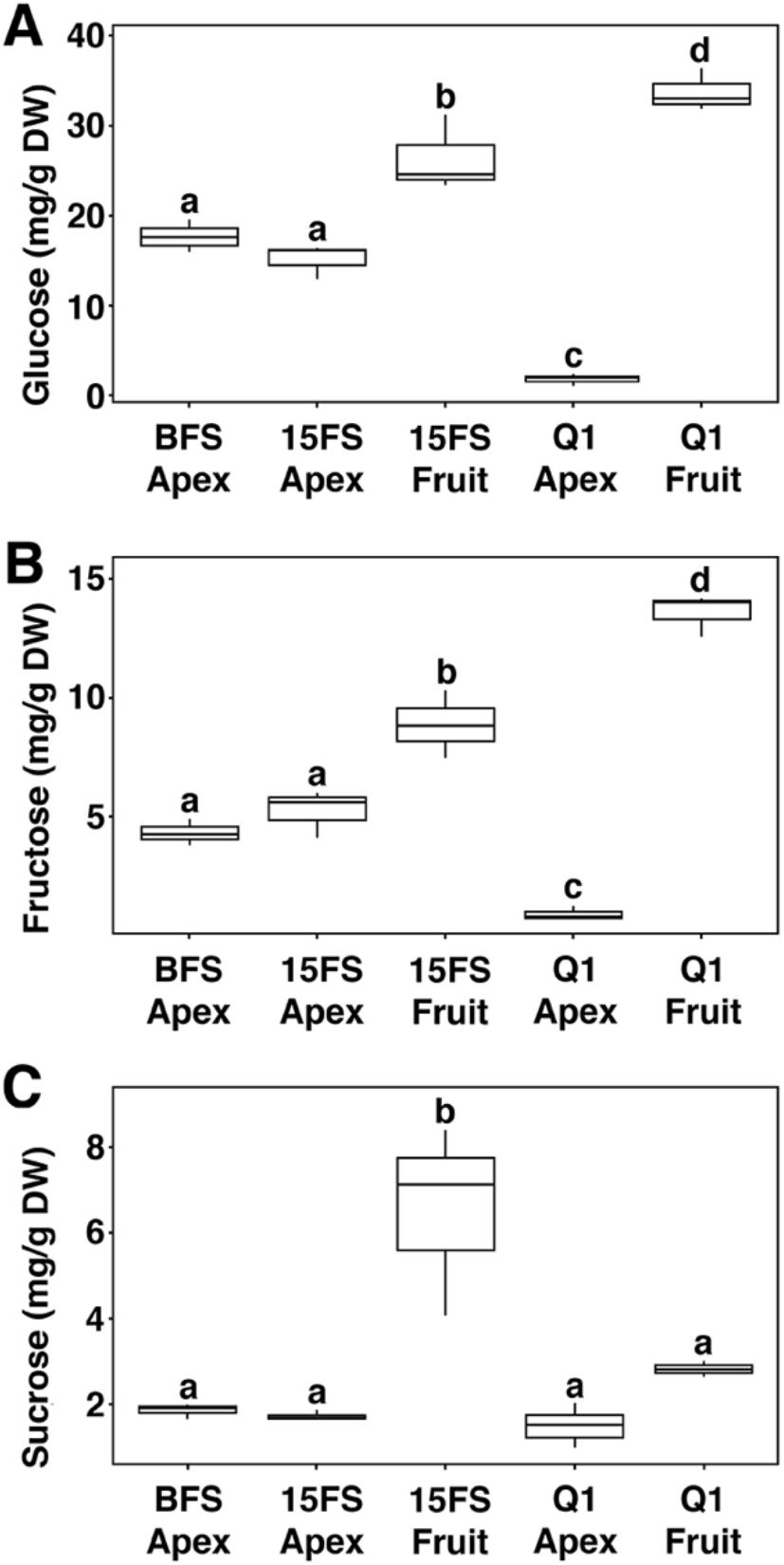
Sugar content in shoot apices and developing fruits. Sugar levels determined in shoot apices before fruit set (BFS Apex), after 15 fruit set (15FS Apex) and at the Q1 stage (Q1 Apex). Sugar levels were also determined in developing fruits when 15 fruits set (15FS Fruit) and at the Q1 stage (Q1 Fruit). The levels of (A) Glucose, (B) fructose and (C) sucrose were measured from dry weight (DW) tissue. The letters above the bars determine whether differences in sugar levels were statistically significant using analysis of variance, Tukey’s honest significant difference.

## DISCUSSION

Dominance interaction between fruits and shoot apices is a major factor that influences shoot architecture and yield (Bangerth, 1989; Smith and Samach, 2013; Walker and Bennett, 2018). During inflorescence development, the decline in meristem size and activity, correlates with a decrease in stem cell renewal based on *WUS* expression (Balanza et al., 2018; Wang et al., 2020). As the growth potential of the inflorescence apex declines, and the meristem become competent for arrest, the end of flowering phase is initiated (Ware et al., 2020). To further extend these studies, our results showed that the end of flowering phase transition is a two-step process. At the Q1 stage, a cessation of growth selectively occurs in the inflorescence apex, while the remaining immature fruits continue to develop. This is supported by expression studies indicating that stem cell renewal and auxin mediated organogenesis in the inflorescence meristem, as well as cell division, are significantly reduced in the Q1 shoot apex. The end of flowering phase is completed at the Q2 stage when fruit growth and development is completed.

Results from a recent study show that production and export of auxin from developing fruits at a late stage of inflorescence development promotes inflorescence arrest (Ware et al., 2020). The end of flowering model proposed by Ware et al., 2020 predicts that auxin export from developing fruits induces inflorescence arrest by disrupting polar auxin transport in the inflorescence stem. Results from our study show that growing fruits impact auxin transport from the inflorescence apex. First, auxin transport in the apical stem was the highest before fruit set. However, after 10 fruits were produced, a significant decline in auxin transport occurred in AS and BS segments. Second, after 20 fruits were produced, an apparent gradual decrease in auxin transport primarily occurred in AS segments. At the Q1 stage, the inhibition of inflorescence growth correlates with a selective dampening of auxin transport in the apical region of the stem below the shoot apex, while transport below the ZFD is functional. Furthermore, an increase in auxin response in the vascular tissues in the ZFD is initiated at Q1 and reaches a maximum at Q2. Finally, removal of fruits at the Q2 stage restores growth and auxin transport in the apical region of the stem. Taken together, we propose that dampening of auxin transport together with an increase in auxin response functions to maintain growth cessation by preventing auxin canalization from the inflorescence apex until the seeds in the developing fruits fully mature.

Sugar signaling plays an essential role in plant developmental phase transitions (Bolouri Moghaddam and Van den Ende, 2013; Poethig, 2013). Flowering is a major phase transition that is regulated by florigen, *FLOWERING LOCUS T* (*FT*), and the age-dependent pathway mediated by the *microRNA156* (*miR156*)/ *SQUAMOSA PROMOTER BINDING PROTEIN-LIKE* (*SPL*) module (Turck et al., 2008; Srikanth and Schmid, 2011; Wang, 2014). Experimental studies show that T6P pathway regulates *FT* and the *miR156*/*SPL* module in leaves and shoot apices, respectively (Wahl et al., 2013). In addition, the suppression of *miR156* during the vegetative phase transition is mediated by sugars, including glucose and sucrose (Yang et al., 2013; Yu et al., 2013), as well as the T6P pathway (Ponnu et al., 2020). In our study, we show that glucose and fructose levels, as well as *TSP1* expression, are significantly reduced in Q1 apices compared to actively growing inflorescence apices. As the biosynthesis of T6P is dependent upon glucose and *TSP1* expression, our results indicate that the T6P pathway is highly reduced in Q1 apices. Therefore, we propose that the suppression of sugar signaling mediated by glucose and the T6P pathway is involved in the end of flowering phase transition.

The reduction in glucose and fructose but not sucrose in Q1 apices at the end of flowering indicates that sucrose metabolism is selectively inhibited in inflorescence apices but not developing fruits. Consistent with this view, *CINV1* expression is suppressed when inflorescence apices transition to the Q1 stage. In contrast, developing fruits at Q1 stage display an increase in sucrose metabolism compared to growing fruits at an earlier stage of inflorescence development. This is supported by the fact that the levels of glucose and fructose are higher, while sucrose is lower in developing fruits at the Q1 stage. Therefore, it is tempting to speculate that end of flowering phase transition not only functions to repress inflorescence growth, but also acts to further increase the growth potential of fruits. The increase in the growth potential of developing fruit at the Q1 stage may be mediated by the end of flowering-competence factors.

Expanding on the model proposed by Ware et al., 2020, we propose that inflorescence growth is maintained by: (1) the basipetal auxin transport system in the apical domain of the inflorescence stem and (2) sugar signaling and metabolism in the inflorescence apex. During the active stage of inflorescence development, the shoot system can support both shoot and fruit growth. However, with the continuous decline in meristem activity and auxin transport out of the inflorescence apex, a competence juncture is reached in which the apical bud including the inflorescence and floral meristems, immature flowers and subtending internodes, can no longer maintain growth, as fruits continue to develop. At this competence juncture, export of auxin from developing fruits selectively impairs auxin transport in the apical stem below the inflorescence apex, which induces inflorescence arrest. We propose that impairment of auxin transport in the apical inflorescence stem suppresses sugar signaling and metabolism in the inflorescence apex, which negatively impacts stem-cell renewal and organogenesis. This is supported by the fact that auxin mediated organogenesis and stem cell renewal is dependent upon sugar availability and signaling (Lauxmann et al., 2016; Pfeiffer et al., 2016). Further, the increase in auxin response in the vascular tissues of the apical stem from the Q1 to the Q2 stage acts to maintain growth arrest by dampening auxin export from the inflorescence meristem and immature floral buds at the Q2 stage.

In fruit tree crops, inhibition of shoot growth by fruit load is a major driver of biennial or alternate bearing, which is a major challenge for fruit tree crop industries worldwide (Samach and Smith, 2013; Smith and Samach, 2013). Due to significant challenges and barriers associated with the usage of genetic and molecular manipulations in fruit tree crops, Arabidopsis may serve as a model system to understand the physiological basis of shoot growth arrest in response to fruit load. Translational research from Arabidopsis to fruit tree crops can be utilized to develop new innovative tools to limit the impact of fruits on shoot growth in order to maximize yield and reduce seasonal variation.

## MATERIALS AND METHODS

### Plant materials and growth conditions

The Arabidopsis thaliana Columbia-0 (Col-0) accession was used to characterize inflorescence arrest in response to fruit load. Auxin response was evaluated during inflorescence development using the *DR5:GUS* system (Ulmasov et al., 1997). Plants were grown at 22°C under long day growth conditions, 16-hour light/8-hour dark.

### Internode measurements

The length of the last 30 internodes were measured in the primary inflorescence after the end of flowering transition was completed in 30 plants. The average length in mm and standard deviation for each internode was determined.

### Gene expression analyses

To examine the expression pattern of key genes that regulate meristem activity and sugar signaling, in situ hybridization was performed using a standard method of fixation, sectioning and mRNA hybridization as previously described (Jackson, 2001; Chuck et al., 2002). Active inflorescence apices were harvested after 5-10 fruits were produced. Quiescent apices were harvested at the Q1 stage of the end of flowering transition. Synthesis of UTP-digoxigenin anti-sense probes were previously described for *STM* (Long et al., 1996), *WUS* (Yadav et al., 2009), *MP* (Zhao et al., 2010) and *TPS1* (Zhao et al., 2010). Primer sequences were used to PCR amplify *CDKB1;1*, *CINV1* and *SUC2* DNA fragments for the synthesis of UTP-digoxigenin antisense probes. The sequences for *CDKB1;1*, *CINV1* and *SUC2* primers were CDKB1;1-F (CGAGATGGACGAAGAAGGTATTCCACC), CDKB1;1-R (<u>GAAATAATACGACTCACTATAGGGACT</u>CGTGAGAAGATCAACTCCTTGAGGTG), CINV1-F (CCGATGGAGATGGCAGAGAGG), CINV1-R (<u>GAAATAATACGACTCACTATAGGGACT</u>GGCCAAGACGCAGATCGCTTGATGAC), SUC2-F (CTGAGTCATGCGATCTCTACTGCG) and SUC2-R (<u>GAAATAATACGACTCACTATAGGGACT</u>CTTACCGCTGCCGCAATCGCTCC). The method to visualize GUS activity in *DR5:GUS* inflorescence shoots was described previously (Sundaresan et al., 1995; Springer et al., 2000). *DR5:GUS* was evaluated in inflorescences during the active period of inflorescence development when apices produced 10-20 fruits, as well as the Q1 and Q2 stages.

### ^14^C-IAA transport assay

Basipetal auxin transport was measured in inflorescence stems using a ^14^C-IAA protocol previously described (Lewis and Muday, 2009). Briefly, 20 mm stem segments were harvested from inflorescences before fruit set. After the shoot apex produced 10, 20 and 35 fruits and at the Q1 and Q2 stages of development, AS and BS segments were isolated from each inflorescence as described in the results section. To measure auxin transport in each stem segment, the apical end of the stem was placed in 20 μL of auxin transport buffer (100 nM ^14^C-IAA, 0.05% MES, pH 5.7). To measure movement mediated by diffusion, a separate set of stem segments were isolated and the apical end of each stem was placed in auxin transport buffer containing naphthylphthalamic acid (NPA) to a final concentration of 10 μM. After 10 hours of auxin transport at 22°C, each segment was removed from the auxin transport buffer (+/-NPA) and a 5 mm section at the apical end was cleaved and discarded. Next, each stem segment was transferred to an Eppendorf tube and ground in scintillation fluid using a plastic pestle. For each sample, the extract from was transferred to a single scintillation vial containing 20 mL of scintillation fluid and ^14^C d.p.m. was determined before conversion to fmoles. Five biological replicates were used to calculate the mean and standard deviation. Analysis of variance and Tukey’s honest significant difference analysis was performed using standard statistical packages in R.

### Sugar measurements

For sugar extraction, ~100 mg of inflorescence apices and developing fruit were collected during inflorescence development in triplicate. After collection, the material was freeze-dried for 12 h and the dry weight for each sample was determined. Each sample was ground with a mortar and pestle in 1.0 mL of 80% ethanol. Next, samples were incubated at 80°C for 30 minutes to extract soluble sugars. After the insoluble material was pelleted at 10,000 xg and the supernatant decanted, the tissues were re-extracted two more times with 80% ethanol. After combining and mixing the three separate 80% ethanol extracts, 650 mL of the soluble extract was placed in an Eppendorf tube and dried in a Gene miVac Quattro (SP Industries, Warminster, PA, USA) for 1.5 h at 55°C. Each dried sample was resuspended in 20 μL sterile H_2_O. Glucose, fructose and sucrose were separated by High Performance Liquid Chromatography using the Sugar-Pak cation-exchange column (Waters, Rydalmere, NSW, AUS). The Aglient Technologies 1200 G1362A infinity refractive index detector (Santa Clara, California, USA) was used to identify and quantify separated sugars in each of the samples by comparison to the glucose, fructose and sucrose standards. Analysis of variance and Tukey’s honest significant difference analysis was performed using standard statistical packages in R.

## ACKNOWLEDGMENTS

We thank Tom Bennett (University of Leeds, UK) for discussions and reviewing the manuscript. We also thank Kate Tepper and Rhys Webber for the maintenance of Arabidopsis plants and Dr Tom Guilfoyle for providing *DR5:GUS* seed.

